# Structural insights into DNA sequence recognition by *Arabidopsis* ETHYLENE RESPONSE FACTOR 96

**DOI:** 10.1101/302299

**Authors:** Chun-Yen Chen, Pei-Hsuan Lin, Kun-Hung Chen, Yi-Sheng Cheng

## Abstract

The phytohormone ethylene is widely involved in many developmental processes and is a crucial regulator of defense responses against biotic and abiotic stresses in plants. Ethylene-responsive element binding protein (EREBP), a member of the APETALA2/ethylene response factor (AP2/ERF) superfamily, is a transcription factor that regulates stress-responsive genes by recognizing a specific *cis*-acting element of target DNA. A previous study showed only the NMR structure of the AP2/ERF domain of AtERF100 in complex with a GCC box DNA motif. In this report, we determined the crystal structure of AtERF96 in complex with a GCC box at atomic resolution. We analyzed the binding residues of the conserved AP2/ERF domain in the DNA recognition sequence. In addition to the AP2/ERF domain, an N-terminal α-helix of AtERF96 participates in DNA interaction in the flanking region. We also demonstrated the structure of AtERF96 EDLL motif, a unique conserved motif in the group IX of AP2/ERF family, is critical for the transactivation of defense-related genes. Our study establishes the structural basis of the AtERF96 transcription factor in complex with the GCC box, as well as the DNA binding mechanisms of the N-terminal α-helix and AP2/ERF domain.

## INTRODUCTION

Plants are exposed to natural environments that may negatively affect their growth and development. To rapidly adapt to environmental change, numerous genes are regulated in response to biotic and abiotic stresses, such as herbivore damage, pathogenic infection, UV irradiation, temperature variation, drought, and high salt content (Ecker, 1995, Kazan, 2015, Mizoi, Shinozaki et al., 2012b, Penninckx, Eggermont et al., 1996, Penninckx, Thomma et al., 1998). These stresses can induce the biosynthesis of ethylene, a gaseous plant hormone confirmed as a mediator of plant stress responses (Ecker, 1995). A *cis*-acting element GCC box motif that constitutes the conserved sequence AGCCGCC is widely present in the promoter region of ethylene-inducible genes and has been suggested as a core sequence of the ethylene-responsive element (ERE) for ethylene signaling in plants (Buttner & Singh, 1997, Hao, Ohme-Takagi et al., 1998, Sessa, Meller et al., 1995, Xu, Narasimhan et al., 1998).

Such a group of transcription factors in plants is activated by ethylene signaling under biotic or abiotic stresses. APETALA2/ETHYLENE RESPONSE FACTORS (AP2/ERFs) are a superfamily of plant-exclusive transcription factors involved in the ethylene-inducible response (Gutterson & Reuber, 2004, Mizoi et al., 2012b). APETALA2 was first isolated from the floral development-related proteins in *Arabidopsis* (Jofuku, den Boer et al., 1994). Additionally, an AP2-LIKE ERE BINDING FACTOR (ERF), also known as ERE BINDING PROTEIN (EREBP), was first isolated as the GCC box-binding protein from tobacco (Ohme-Takagi & Shinshi, 1995). AP2/ERFs contain a highly conserved DNA-binding domain consisting of approximately 60 amino acids (Nakano, Suzuki et al., 2006), specifically to recognize the GCC box at the upstream operators of defense-related genes, such as pathogen-inducible *PLANT DEFENSIN 1.2* (PDF1.2) and *PATHOGENESIS*-*RELATED PROTEIN 4* (*PR4*) (Ohme-Takagi & Shinshi, 1995, Shinshi, Usami et al., 1995). Therefore, AP2/ERFs play an important role in the regulation of defense responses in plants.

The AP2/ERF family has 147 members that can be divided into three subfamilies: AP2, RAV, and ERF (Nakano et al., 2006). The AP2 subfamily is composed of one or two AP2 domains, which primarily participate in the process of floral development (Elliott, Betzner et al., 1996). The RAV subfamily contains an ERF domain and a B3 DNA-binding domain involved in ethylene and brassinosteroid responses (Hu, Wang et al., 2004). The largest ERF subfamily comprises 122 members and mainly involves an ERF domain that interacts with the GCC box DNA sequence (Hao et al., 1998). The ERF family can also recognize non-GCC box *cis* elements. For instance, the DEHYDRATION-RESPONSIVE ELEMENT BINDING PROTEINS (DREB1A and DREB2A) bind to the DRE box (5’-[A/G]CCGAC-3’) of dehydration- and cold stress-related genes (Liu, Kasuga et al., 1998). AtERF13, RAP2.4, and RAP2.4L are involved in the abscisic acid (ABA) response by binding to the ABA-related cis-acting coupling element (CE1) upstream of the CE1 BINDING FACTOR (CEBF) in *Arabidopsis* (Lee, Park et al., 2010, Novillo, Medina et al., 2007). The ERF subfamily can be further divided into ten groups (groups I–X) based on sequence similarity. Among these, groups VIII and IX play an important role in the interaction between plant and pathogen (Nakano et al., 2006). The members of group IX-c, such as OCTADECANOID-RESPONSIVE AP2/ERF 59 (ORA59), TRANSCRIPTIONAL REGULATOR OF DEFENSE RESPONSE 1 (TDR1), AtERF14, AtERF15, as well as members of group IX-a, such as AtERF1 and AtERF13, positively regulate pathogen defense responses by recognizing the GCC box of defense-related genes basic chitinase (*Chi*-*B*) and *PDF1.2* (Berrocal-Lobo, Molina et al., 2002, Gutterson & Reuber, 2004, Onate-Sanchez, Anderson et al., 2007, Pre, Atallah et al., 2008, Zhang, Huang et al., 2015). Furthermore, group IX-c members of the AP2/ERF family interact with MEDIATOR25 (MED25) using a highly conserved sequence EDLL motif (also known as CMIX-1) at the C-terminus (Nakano et al., 2006, Tiwari, Belachew et al., 2012). Yeast two-hybrid analysis revealed that AtERF98 loses its binding function to MED25 when the conserved leucine is replaced by valine in the EDLL motif (Tiwari et al., 2012).

*Arabidopsis* ETHYLENE RESPONSE FACTOR 96 (AtERF96, AT5G43410), a member of ERF group IX, contains an AP2/ERF domain and an EDLL motif, and positively regulates defense responses against the necrotrophic pathogens *Botrytis cinerea* and *Pectobacterium carotovorum* (Catinot, Huang et al., 2015). Previous studies indicate that the expression level of AtERF96 can be induced via ethylene and jasmonate (JA) signaling pathways, but is antagonized by the salicylic acid (SA) response (Catinot et al., 2015, Zander, Thurow et al., 2014). The impact of TGACG SEQUENCE-SPECIFIC BINDING PROTEINS (TGA2, TGA5, TGA6) on AtERF96 mRNA levels has been confirmed (Zander et al., 2014). Additionally, microarray analysis revealed that 45% of the 126 up-regulated genes in transgenic overexpressing AtERF96 are strongly related to the plant defense response (Catinot et al., 2015).

To date, we lack a full-length protein structure of the AP2/ERF transcription factor. However, the GCC box-binding domain (GBD, namely, the AP2/ERF domain) of AtERF100 (previously designated as ERF1 and renamed At4g17500 by Nakano *et al.*, 2006) was previously determined using nuclear magnetic resonance spectroscopy (NMR; PDB ID: 1GCC) (Allen, Yamasaki et al., 1998). The AP2/ERF domain is composed of three β-sheets and an α-helix; the β-sheets interact monomerically with the target 11 base pairs of double-strand DNA (5’-GCTAGCCGCCAGC-3’). Even if the classification of AP2/ERFs provides some information on their potential role in plants, only a limited number of AP2/ERF transcription factors have been functionally characterized. Here, we solved the crystal structure of the AtERF96–DNA complex, including an AP2/ERF domain and a unique EDLL motif. We determined the AtERF96–GCC11 binding mechanism, demonstrated that the N-terminal α-helix of AtERF96 binds to the DNA minor groove, and clarified the influence of AtERF96 on the target gene expression.

## MATERIALS AND METHODS

### Cloning, expression, and purification of AtERF96

The cDNA of full-length *Arabidopsis thaliana ERF96* (AT5G43410) was cloned into pET28a expression vector (Novagen) with a hexahistidine tag (6xHis-tag) at the N-terminus. The expression vector was transformed into *Escherichia coli* strain BL21 (DE3) (Novagen), and then incubated at 37°C in a 2 L flask with shaking until 0.4–0.6 absorbance at 600 nm was achieved. The expression of AtERF96 protein was induced by 0.1 mM isopropyl β-D-1-thiogalactopyranoside (IPTG) for 18 h at 16°C. The cells were harvested by centrifugation at 9,820 *g* for 30 min at 4°C, then resuspended and lysed in lysis buffer (30 mM HEPES pH 8.0, 500 mM NaCl, 20 mM imidazole, 0.5 mg/mL DNase I). The cells were lysed by sonication and the cell debris was removed via centrifugation at 18,900 *g* for 25 min at 4°C. Protein purification was performed by Fast Protein Liquid Chromatography (FPLC) using the AKTA prime plus system (GE Healthcare). The filtered supernatant was applied to a 5 mL HisTrap™FF column (GE Healthcare) pre-equilibrated with binding buffer (30 mM HEPES pH 8.0, 500 mM NaCl, 20 mM imidazole). Proteins were eluted with elution buffer (30 mM HEPES pH 8.0, 500 mM NaCl, 500 mM imidazole). The purified protein solution was applied to a HiTrap Heparin HP column (GE Healthcare) to remove endogenous DNA fragments of the host cells. The column was pre-equilibrated with buffer A (30 mM HEPES pH 8.0), and the protein was eluted using the linear gradient of buffer exchange from 0 to 100% with buffer B (30 mM HEPES pH 8.0, 1 M NaCl). The eluted proteins were further desalted to storage buffer (30 mM HEPES pH 8.0, 250 mM NaCl, 10% glycerol) using a HiTrap™ Desalting column (GE Healthcare). Purified protein concentrations were determined at 595 nm absorbance using an ELISA reader (FlexStation 3, Molecular Devices) and the standard curve method. The Coomassie brilliant blue G250 (CBB) protein assay solution (5×) (Bio-Rad) was used as the blank solution, and a dilution series of CBB mixed with 5 μL bovine serum albumin (BSA) at 1000, 500, 250, and 125 μg/mL were used as standards. The purified AtERF96 proteins were concentrated to 10 mg/ml using 10 kDa centrifuge tubes (Amicon Ultra-15 Centrifugal Filter; Merck Millipore) and stored at –80°C.

### Site-directed mutagenesis

The *AtERF96* single and double mutations were generated using the QuikChange Lightning Site-Directed Mutagenesis Kit (Agilent). The *AtERF96* mutations were introduced into cDNA fragments through PCR using the primers listed in Table S1, and the fragments were cloned into the pET28a expression vector. Protein expression and purification were performed as for wild-type ERF96.

### Preparation of fluorescein-labelled double-stranded DNA probes

Individual single-stranded oligonucleotides of GCC-box fragments were synthesized by commercial gene synthesis (Genomics). The 5’ end of forwarding oligonucleotides were labeled with fluorescein. Annealing of the two complementary strands was performed in 30 mM HEPES (pH 7.4) by heating at 95°C for 1 min, followed by slowly cooling to room temperature for 20 min. The concentration of the annealed DNA probes was measured by spectrophotometry (DS-11, DeNoVix) and stored at –20°C.

### Size-exclusion chromatography (SEC)

The SEC column was pre-equilibrated with one column volume of GF1 buffer (30 mM HEPES pH 8.0, 500 mM NaCl, 10% glycerol). To determine the molecular weight of the target protein, 300 μL of protein standard (Bio-Rad) containing γ-globulin (bovine, 158 kDa), ovalbumin (chicken, 44 kDa), myoglobin (horse, 17 kDa), and vitamin B12 (1.35 kDa) was applied to the column, and the elution volumes of the standards were plotted against the logarithm of the standards’ molecular weights.

The polymer characterization of AtERF96 proteins was performed by the AKTA prime plus system (GE Healthcare) using a Superdex 75 column (GE Healthcare). The AtERF96 protein sample (< 5 mL volume) was applied to the column, and the fractions of elution peaks, including monomeric AtERF96 proteins, were pooled and concentrated to 10 mg/ml using a 10 kDa centrifuge tube.

The binding analysis of ERF96–GCC-box was performed by the AKTA prime plus system (GE Healthcare) using a Superdex 75 column (GE Healthcare). A mixture of AtERF96 protein and double-stranded GCC12 DNA fragments were pre-incubated on ice at a 1:2 molar ratio for 30 min. The AtERF96 protein, double-stranded GCC12 DNA fragments, and the ERF96–GCC12 complex were applied to the column with GF2 buffer (30 mM HEPES pH 8.0, 62.5 mM NaCl, 10% glycerol). The fractions of each elution peak were pooled and concentrated to 10 mg/ml using a 10 kDa centrifuge tube.

### Fluorescein-based electrophoretic mobility shift assay (fEMSA)

The fluorescein-labeled probes and AtERF96 recombinant proteins were incubated in 30 mM HEPES (pH 7.4), 62.5 mM NaCl, and 5% glycerol for 30 min at room temperature in the dark. A 10% polyacrylamide gel was pre-run at 120 V for 40 min at 4°C. Samples were mixed with 5× loading dye and run at 120 V for 90 min at 4°C in the dark. The gel was scanned for fluorescent band shift using the FluorChem™ M system (ProteinSimple) and a luminescence imaging system (Fuji LAS-3000). Quantitative analysis of fluorescent band shift was performed using ImageJ software, with the band intensity of the AtERF96-GCC box set as a baseline (100%) to determine the relative binding level.

### Dynamic light scattering (DLS) assay

The purified AtERF96 protein sample (20 μL at 1 mg/mL) was loaded into the cuvette, and the particles of the protein molecules in the solutions were measured at 25°C. The batch light scattering data were recorded using the DynaPro Plate Reader I (Wyatt Technology) and analyzed with DYNAMICS 7.0 software (Wyatt Technology). The diffusion coefficient was calculated from the intensities of light scatter from the molecule particles, and further analysis of the hydrodynamic radius, diameters, and molecular weights of the target protein particles by the Stokes-Einstein Law is described as follows:

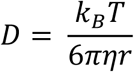

where D is the diffusion constant (m^2^/s), k_B_ is Boltzmann constant, T is the absolute temperature, *π* is the ratio of a circle’s circumference to its diameter, *η* is the dynamic viscosity, and *r* is the radius of the spherical particle.

### Fluorescence polarization (FP) assay

Purified AtERF96 proteins were desalted to FP buffer (30 mM HEPES pH 7.4, 250 mM NaCl, 10% glycerol) using a HiTrap™ Desalting column (GE Healthcare), and the concentration was determined by the Bradford protein assay. The AtERF96 protein samples were two-fold serially diluted in FP butter to 16 or 24 concentrations, and 50 μL of each diluted protein sample was added to a 96-well plate, along with 50 μL of 20 nM GCC12 probes in each sample well. A set of wells containing 100 μL FP buffer, and another set of wells containing 50 μL FP buffer mixed with 50 μL of 20 nM GCC12 probes were used as blanks and controls, respectively. FP enables the study of molecular interactions by monitoring changes in the apparent size of fluorescently-labelled or inherently fluorescent molecules, which are often referred to as the tracer or the ligand (Checovich, Bolger et al., 1995, Heyduk, Ma et al., 1996, Moerke, 2009). The samples of fluorescent molecules were excited by plane-polarized light, and the emission spectra were recorded and analyzed by PARADIGM™ (Beckman Coulter/Molecular Devices). Quantification of fluorescence polarization (FP) is defined as the difference between the emission intensities of horizontally (*F*_∥_) and perpendicularly polarized light (*F*_⊥_) to the excitation light plane normalized by the total fluorescence emission intensity (Moerke, 2009). The formula of FP is described as follows:

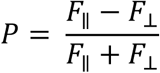

where P is the polarization obtained by subtracting the blank value of both the horizontally and perpendicularly polarized light. The anisotropic levels of polarized fluorescence were plotted against the concentrations of protein samples using Prism 7 (GraphPad Software, Inc.) with the two-site binding equation. The dissociation constant (K_d_) is determined by the correlation between polarizations and sample concentrations, and the formula of two-site binding is described as follows:

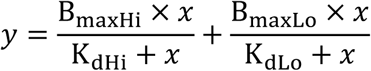

where *x* is the protein concentration, *y* is the polarized value. B_maxHi_ and B_maxLo_ are the maximum specific bindings to the two sites in the same units as *y*. K_dHi_ and K_dLo_ are the equilibrium binding constants, in the same units as *x*.

### Protoplast transactivation analysis (PTA)

The experimental procedure is described in a previous report (Yoo, Cho et al., 2007). Wild-type *Arabidopsis* plants were grown on sterile soil in an environment-controlled chamber. True leaves number 5–7 from 4-week-old plants were chosen before flowering and 1-mm leaf strips were cut from the middle part of a leaf. Leaf strips were quickly and gently transferred into the prepared enzyme solution (20 mM 2-(N-morpholino) ethanesulfonic acid (MES) pH 5.7, 1.5% (w/v) cellulase R10, 0.4% (w/v) macerozyme R10, 0.4 M mannitol, 20 mM KCl). To enhance enzyme solubility, the solution was heated at 55°C for 10 min to inactivate DNase and proteases. While the solution was cooling to room temperature (25°C), 10 mM CaCl_2_, 1 mM β-mercaptoethanol, and 0.1% BSA were added. Leaf strips were vacuum-infiltrated for 30 min in the dark using a desiccator, then the digestion was continued in the dark at room temperature for at least 3 h. The enzyme/protoplast solution was diluted with an equal volume of W5 solution (2 mM MES pH 5.7, 154 mM NaCl, 125 mM CaCl_2_, 5 mM KCl) before filtration to remove undigested leaf tissues. The enzyme/protoplast solution was filtered using 75-μm nylon mesh wetted with W5 solution, centrifuged at 200 *g* for 2 min to pellet the protoplasts, then the supernatant was removed and the pellet was re-suspended in W5 solution with gentle swirling. Protoplasts were centrifuged again for 15 min to remove W5 solution, and the protoplast pellet was re-suspended with MMG solution (4 mM MES pH 5.7, 0.4 M mannitol, 15 mM MgCl_2_) at room temperature. Ten μl of DNA plasmids and 100 μl of protoplasts were gently mixed in the microfuge tube. Then, 110 μl polyethylene glycol (PEG) solution (40% (w/v) PEG4000, 0.2 M mannitol, 100 mM CaCl_2_) was added and mixed gently, and the transfection mixture was incubated at room temperature for 10 min. The transfection was stopped by diluting the mixture with 400 μl W5 solution and gentle mixing. The protoplast mixture was centrifuged at 100 *g* for 2 min to remove the supernatant and re-suspended gently with 1 ml WI solution (4 mM MES pH 5.7, 0.5 M mannitol, 20 mM KCl). Protoplasts were transferred to a tissue culture plate and incubated at room temperature for 8 h, then re-suspended and harvested by centrifugation at 100 *g* for 2 min to remove the supernatant. Protoplast lysis buffer (100 μl) was added to the protoplasts and mixed vigorously by vortexing for 10 s, then incubated on ice for 5 min and centrifuged at 1000 *g* for 2 min. Twenty μl of lysate was added to 100 μl luciferase mix (Dual-Luciferase^®^ Reporter Assay System, Promega), and the luciferase activity was measured with a luminometer (Infinite M200 pro, TECAN).

### Protein crystallization and data collection

The AtERF96 protein and GCC11 double-stranded DNA probe (5’-TAGCCGCCAGC-3’) were incubated in a tube at a 1:2 molar ratio, and concentrated to 6 mg/mL with GF2 buffer for crystallization. Screening for suitable crystallization conditions was performed using the Crystal Phoenix Liquid Handling System robot (Art Robbins Instruments, LLC). The program was set to a sitting-drop method, which dispensed an equal volume of the protein–DNA mixture and screening buffer to a volume of 1 μL to each well of a 96-well plate. The AtERF96–GCC11 complex crystals were observed at a temperature of 295 K at four crystallization conditions: Natrix™ No. 45 (0.05 M Tris-HCl pH 8.5, 0.025 M MgSO_4_·H_2_O, 1.8 M (NH_4_)_2_SO_4_), Natrix™ 2 No. 6 (0.05 M sodium cacodylate trihydrate pH 6.0, 35% tacsimate pH 6.0), Natrix™ 2 No. 26 (0.05 M 3-(N-morpholino)propanesulfonic acid (MOPS) pH 7.0, 0.02 M MgCl_2_·6H_2_O, 55% tacsimate pH 7.0), and PEGRx™ 2 No. 6 (0.1 M sodium citrate tribasic dihydrate pH 5.0, 10% (v/v) 2-propanol, 26% (v/v) PEG 400; Hampton Research, Inc.). Crystals grew to a suitable size for X-ray diffraction after six months. All diffraction data were collected at 100 K on beamline 13C1 at the National Synchrotron Radiation Research Center (NSRRC), Hsinchu, Taiwan. Diffraction data were recorded using the ADSC Quantum-315r CCD detector and collected using Blu-Ice software (McPhillips, McPhillips et al., 2002).

### Structure determination and refinement

Diffraction data were indexed, integrated, and scaled using the HKL2000 package (Otwinowski & Minor, 1997). The crystallographic structure was solved by the PHENIX platform (Adams, Afonine et al., 2010). The assessment of data quality was analyzed by the phenix.xtriage program. Data from the crystal that grew in Natrix™ 2 No. 6 had the best diffraction quality, and the resolution limit reached 1.76 Å. The AtERF96–GCC11 complex was co-crystallized in space group P1 2_1_ 1, which comprised of one AtERF96 and one GCC11 DNA fragment in an asymmetric unit. Twinning analysis by the phenix.xtriage program showed that the data consists of five pseudo-merohedral twins with 3-fold axes (-h-l, k, h/ l, k, -h-l) and 2-fold axes (l, -k, h/ h, -k, -h-l/ -h-l, -k, l). The structure of the AtERF96–GCC11 complex was solved by the molecular replacement method with Phaser (McCoy, Grosse-Kunstleve et al., 2007), using the structure of the AtERF100 AP2/ERF domain (K144-V206) (Protein Data Bank [PDB] ID: 1GCC) (Allen et al., 1998) as a template model. The unknown region of the AtERF96 structure was built manually using COOT software, according to the *F*_o_–*F*_c_ electron density map. The resulting electron density map was sharpened by density modification using RESOLVE (Afonine, Grosse-Kunstleve et al., 2012). Refinement was continued with several cycles of positional, B-factor, occupancies, and TLS (Translation-Libration-Screw-rotation) refinement. Data was detwinned against the twin operators by phenix.xtriage, and further improvement of the density map was achieved by using the twin fraction refinement by REFMAC5 of the CCP4 platform (Murshudov, Skubak et al., 2011, Winn, Ballard et al., 2011) to filter out those small twin fractions so that the major twin domain remains. The revised structure factor data was refined again by phenix.refine, using new R-free flag for several cycles. Validation was performed by the MolProbity program (Chen, Arendall et al., 2010) to check the real-space correlation, molecular geometry, and Ramachandran plots. The stereochemistry of AtERF96 revealed Ramachandran outliers at 4.7%, including Phe71, Pro72, Val81, Gln103, Val104, and Val114. Many Ramachandran outliers were caused by the weak electron density in disordered regions. All structural models were generated using PyMOL (Schrödinger, LLC).

### Data availability

Atomic coordinates and structure factors for the reported crystal structures have been deposited with the Protein Data Bank under accession number 5WX9.

## RESULTS

### AtERF96 recognizes the core sequence of the GCC box motif

The full-length AtERF96 protein consists of 131 amino acids. We constructed and expressed a series of AtERF96 proteins, including wild-type and different mutants, in an *E. coli* system, and purified these using fast protein liquid chromatography (FPLC). Two elution peaks of the AtERF96 protein from the size-exclusion chromatography (SEC) analysis were determined at approximately 286.5 and 26.7 kDa (Fig. S1A). However, the molecular weight of the AtERF96 protein ranges between 15 and 20 kDa based on SDS-PAGE (Fig. S1B). We used dynamic light scattering (DLS) to further confirm protein homogeneity and the size distribution profile. The results showed that the precise monomeric form (62.7 mL) was 17 kDa, which is consistent with the results of the SDS-PAGE analysis (Fig. S1D and E).

The AtERF family widely regulates defense-related genes by recognizing the GC-rich sequences at the upstream promoter. Hence, we designed the SEC experiments to clarify whether the AtERF96 protein interacts with the GCC box motif. An earlier elution volume of the SEC trace indicated that the AtERF96 protein interacts with the GCC12 DNA probe composed of 12 base pairs (Fig. 1A). To determine whether the length of the GCC box sequence influences the binding ability of AtERF96, we designed different lengths of GCC probes comprised of a core sequence and a variable flanking region according to the GC-rich promoter sequence in *Arabidopsis* (Table S2). Fluorescence-based electrophoretic mobility shift assay (fEMSA) analysis showed that all GCC probes bound to the AtERF96 protein, especially GCC11 and GCC12 (Fig. 1B). Therefore, we co-crystallized AtERF96 with the GCC11 DNA site and determined the structure to a resolution of 1.76 Å with final R_work_/R_free_ values of 20.7%/22.7% (Fig. S1C, Table 1).

**Figure 1.**
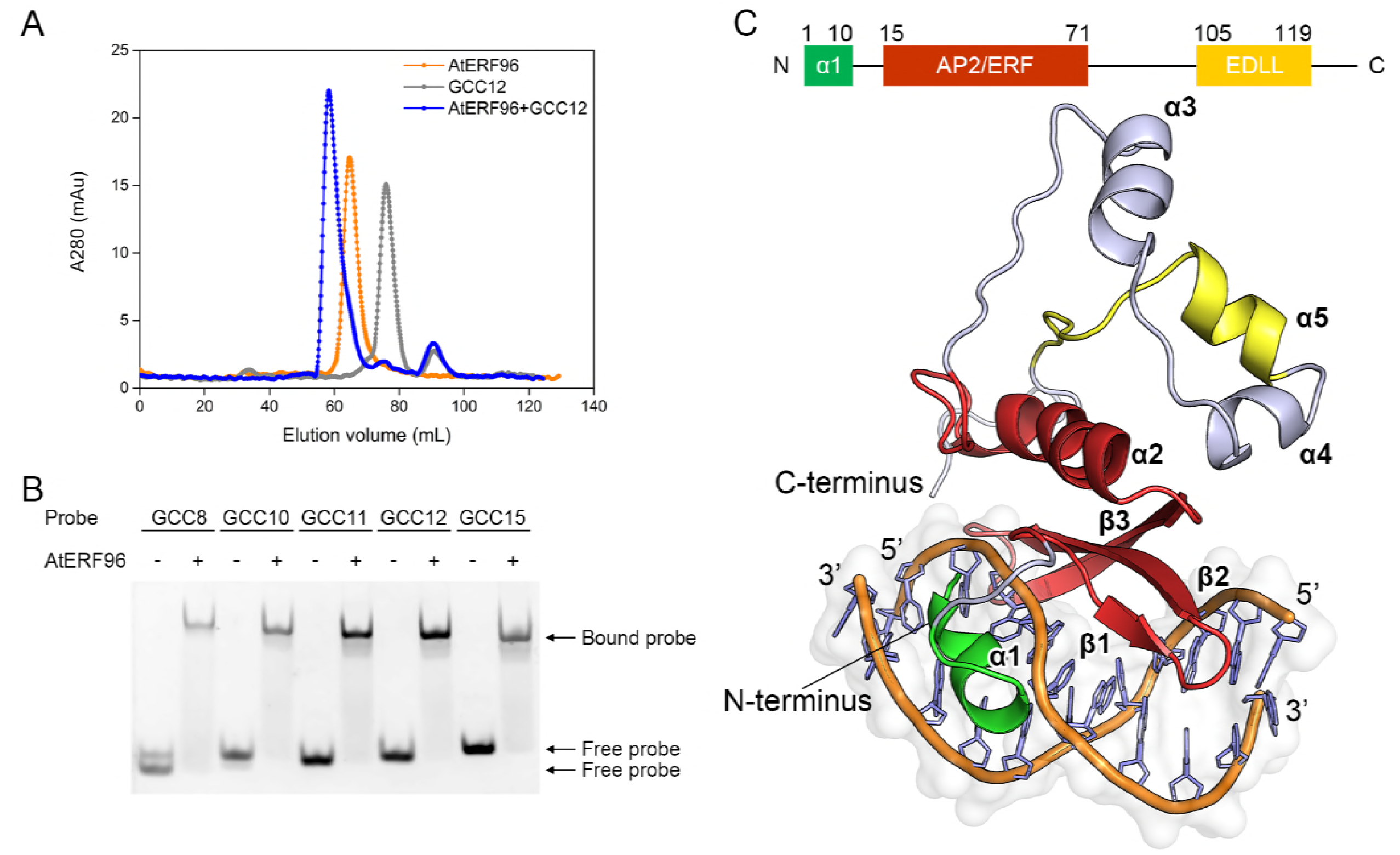
Characterization and crystal structure of the AtERF96–GCC box complex. (**A**) SEC traces of the AtERF96 protein, GCC12 probe, and AtERF96–GCC12 complex are shown as orange, gray, and blue lines. (**B**) EMSA binding analysis of AtERF96 proteins with various GCC probes. (**C**) Ribbon representation of crystal structure of the AtERF96–GCC11 complex. The N-terminal binding region, AP2/ERF domain, and C-terminal EDLL motif are colored in green, red, and yellow.

**Table 1.** Data collection and refinement statistics (molecular replacement)

| AtERF96–GCC-box |  |
| --- | --- |
| **Data collection** |  |
| Space group | P 1 2<sub>1</sub> 1 |
| Beamline | BL13C1, NSRRC |
| Wavelength (Å) | 0.97622 |
| Cell dimensions |  |
| *a*, *b*, *c* (Å) | 39, 81.2, 39 |
| $\alpha$, $\beta$, $\gamma$ (°) | 90, 120, 90 |
| Resolution (Å) | 50-1.76 (1.83-1.76)* |
| Total reflections | 67975 (7034) |
| Unique reflections | 19986 (1986) |
| Mean I/sigma(I) | 10.45 (4.04) |
| Multiplicity | 3.4 (3.5) |
| Completeness (%) | 97.3 (98.9) |
| Wilson B-factor (Å<sup>2</sup>) | 9.30 |
| R-meas (%) | 6.22 |
| R-merge (%) | 5.17 (34.2) |
| CC(1/2) | 99.7 (86.6) |
| **Refinement** |  |
| Resolution (Å) | 21.1-1.76 (1.83-1.76)* |
| R<sub>work</sub>/R<sub>free</sub> (%) | 20.7/22.7 |
| Reflections (work/test) | 17966/2003 |
| No. atoms |  |
| Protein | 1008 |
| DNA | 445 |
| Water | 338 |
| R.m.s deviations |  |
| Bond lengths (Å) | 0.014 |
| Bond angles (°) | 1.772 |
| Ramachandran plot (%) |  |
| Favored, allowed, outliers | 91.5, 3.8, 4.7 |
| B-factor (Å<sup>2</sup>) |  |
| Average | 19.18 |
| Protein | 19.51 |
| DNA | 20.19 |
| Water | 16.87 |

\*Values in parentheses are for the highest-resolution shell.

### Crystal structure of the AtERF96–GCC11 complex

The complex structure consists of the AtERF96 protein with all 131 amino acids, and a double-stranded GCC box motif with 11 base pairs (Fig. 1C). The AtERF96 structure is composed of five α-helices and three β-sheets, including an AP2/ERF domain (K14–E74, β1–β3) for target gene recognition, as well as an EDLL motif (F105–L119, α5) for transcriptional activation. The three-stranded antiparallel β-sheet fragment of the AP2/ERF domain binds to the GCC11 motif and crosses the adjacent major groove region. We found that AtERF96 and AtERF100 could be superimposed with a backbone root-mean-square deviation of 1.31 Å across 55 Cα atoms in the AP2/ERF domain (Fig. S2). The front of the AP2/ERF domain is the N-terminal α-helix (M1–G9, α1), which docks into the minor groove of the GCC11 motif. A linker consisting of eight residues (A10–G17) connects the α1 helix and β1 sheet, gripping one strand of the DNA double helix between the α1 helix and the β1 sheet of the AP2/ERF domain (Fig. 1C). Extending from the AP2/ERF domain, three α-helices (α3–α5), including an EDLL motif, constitute the C-terminal region (Y75–K131) in a triangular-shaped architectural design. Most residues of the AP2/ERF domain have a positively charged electric potential and are highly conserved in group IX of the AP2/ERF family (Fig. 2A, Fig. S3). The AP2/ERF domain consists of several arginines in the three-stranded antiparallel β-sheets bound to the thymines or guanines of the GCC box, which generates a protein-DNA binding network (Fig. 2B, Table S3). Residues R16, R31, and R39 of the AP2/ERF domain interact with the phosphate group of base G15, as well as the guanine group of bases G6, G15, and G16 (Fig. 2C). Residues R19 and R21 interact with bases T1, G3, G18, and G19 of the GCC11 motif at the adjacent interface (Fig. 2D). Furthermore, the conserved tryptophans W23 and W41 provide hydrophobic interactions to stabilize nucleobases T1, G3, and C4 of the GCC11 motif (Fig. 2E). We observed that residues D2, Q3 and R6 of the N-terminal α1 helix bind to nucleotides G10, C11, and T14 of the GCC11 motif and partially disturb the interactions of DNA base pairs in the 3’ flanking region (Fig. 2F). We therefore analyzed the nucleic acid structure of AtERF96–GCC11 using the w3DNA server (Zheng, Lu et al., 2009). The conformational analysis indicated that the GCC11 structure shows an obvious shift and twist in the base step C7/C8, as well as a large tilt and roll in the base step T14/G15 (Table S4). The parameters imply hydrogen bond disruption of DNA base pairs C8-G15 and A9-T14 (Table S5 and S6). The results suggest that the AtERF96 protein specifically binds the GCC box core sequence through the AP2/ERF domain, and also connects the N-terminal α1 helix to these interactions by binding to the 3’ flanking region of GCC11.

**Figure 2.**
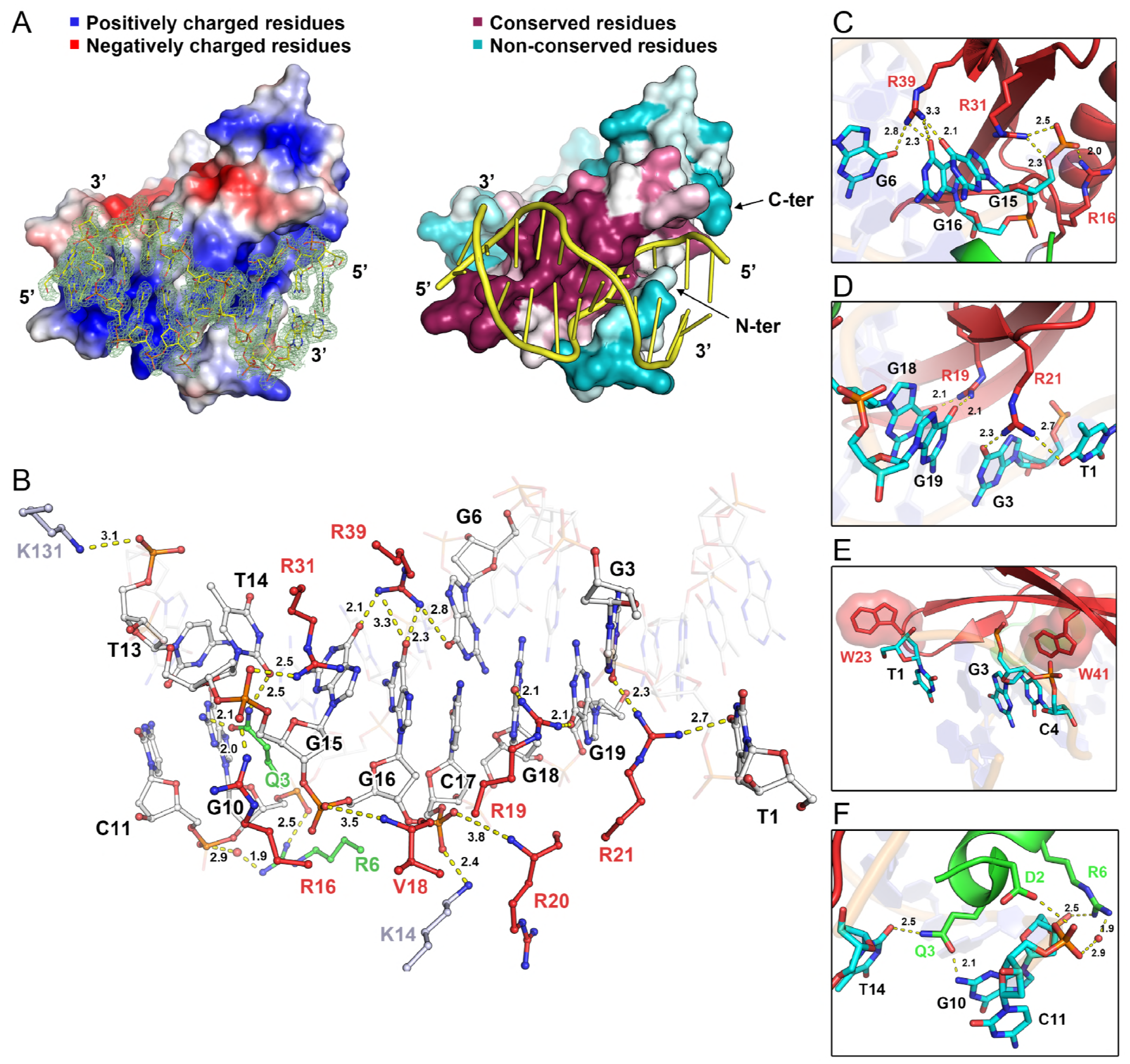
Insights into the interaction of the AtERF96–GCC11 complex. (**A**) Surface representation of electrostatic potential (left) and sequence conservation (right) of the AtERF96–GCC11 complex. The positively and negatively charged residues are indicated as blue and red color on the electrostatic model. The conserved and non-conserved residues are indicated as crimson and blue-green color on the sequence conservation model. (**B**) Zoom-in view of the interaction interface of AtERF96–GCC11 complex. Residues of the AP2/ERF domain, the N-terminal region, and the others interact with DNA are shown as red, green, and grey sticks, respectively. The binding nucleotides are represented as white sticks. (**C-F**) Zoom-in view of the interaction interface of AtERF96-GCC11 complex. (**C**) Ionic interaction of AtERF96 residues R16, R31, R39 with nucleotides G6, G15, G16 are shown as sticks. (**D**) Ionic interaction of AtERF96 residues R19, R21 with nucleotides T1, G3, G18, G19 are shown as sticks. (**E**) Aromatic interaction of AtERF96 residues W23 and W41 with nucleotides T1, G3, and C4. (**F**) Ionic interaction of AtERF96 residues D2, Q3 and R6 with nucleotides G10, C11 and T14. The conserved residues of the AP2/ERF domain are shown as red sticks, and the N-terminal binding residues are shown as green sticks. The binding nucleotides are represented as cyan sticks. Dashed yellow lines indicate a potential interaction network with bond lengths, and water molecules are shown as a red sphere.

### Effect of mutations on the AtERF96–GCC box interaction

In view of the structural information about the AtERF96–GCC box complex, we investigated the importance of conserved residues in the AP2/ERF domain of AtERF96 for GCC box binding. We present a series of AtERF96 mutants corresponding to the binding residues of the structural data and analyzed the dissociation constant (K_d_) with a fluorescently labeled GCC box probe using a fluorescence polarization (FP) assay (Fig. 3A). We chose the GCC12 probe to perform the analysis due to its significant binding shift in the fEMSA assay (Fig. 1B). The results showed that the curves of concentration-dependent polarization fit two sites binding with two independent K_d_ values (K_dHi_ and K_dLo_) (Table S7). AtERF96 mutants had a significant reduction in the binding ability of R16A, R19A, R21A, R39A, and R41A to the GCC12 probes (Fig. 3, Table S7). All of above mutants showed raised levels of K_dHi_ value, implying that these residues are necessary for specific binding in the GCC box. Except for the R16A, W23A, and double-mutant proteins, most mutants remained roughly at the same level of K_dLo_ relative to the wild-type (Fig. 3, Table S7). The raised K_dLo_ levels of R16A and W23A reflected that these residues are involved in non-specific binding in the GCC box, including the π-π stacking of the indole ring and phosphate group binding (Fig. 2C and E). The R19A/R21A and R31A/R39A mutants showed a severe interference in the binding to the GCC12 probes (Fig. 3J and K, Table S7), indicating that these double mutants nearly lost their ability to recognize the core sequence. We noticed that the values of K_dHi_ in the R39A, R19A/R21A, and R31A/R39A mutants were approximately equal to the K_dLo_ values (Fig. 3J and K). Thus, we further analyzed all the polarization data using the equation of one-site binding. The results showed that the polarized curves of R19A, R21A, R39A, W41A and double mutants could be also fitted by the one-site binding (Fig. S4). In addition, R39A, R19A/R21A, and R31A/R39A mutants revealed the similar K_d_ value to the K_dHi_ and K_dLo_, respectively (Fig. S4G, I, and J, Table S7). This indicates that the binding specificity of these mutants was weakened as the features of two-site binding became insignificant. We also performed a fEMSA assay to verify the binding ability of various AtERF96 mutants with different lengths of the GCC box probe. Irrespective of the GCC box probe length used, the binding affinities of the R19A, R21A, R31A, R39A, and W41A mutants were severely decreased (Fig. S5 and S6). The R16A mutant showed minor affinities with the shorter GCC8 and GCC10 probes (Fig. S5A and B), whereas the R19A/R21A and R31A/R39A double mutants barely had the ability to bind any probes (Fig. S7A). Similar to the results of the FP analysis, the W23A mutant showed a lower binding ability with the GCC8, GCC11, and GCC15 probes, and the W41A mutant revealed a more severely reduced interaction with all of the GCC probes (Fig. S5 and S6). We further investigated the importance of AtERF96 N-terminus in GCC box binding, and designed a series of AtERF96 mutants in view of the probable DNA-binding residues in the α1 helix (Fig. 4A). All N-terminal mutants had limited influence on K_dHi_ levels, except for the N-terminal truncated protein (Fig. 4B–F). However, the K_dLo_ levels of D2A/Q3A/R6A and ND10 significantly increased (Fig. 4E and F, Table S7). The results indicate that the residues in the α1 helix are involved in non-specific binding. Overall, most of the conserved arginines and tryptophans in the AP2/ERF domain of the AtERF96 protein are crucial for recognizing the GCC box motif.

**Figure 3.**
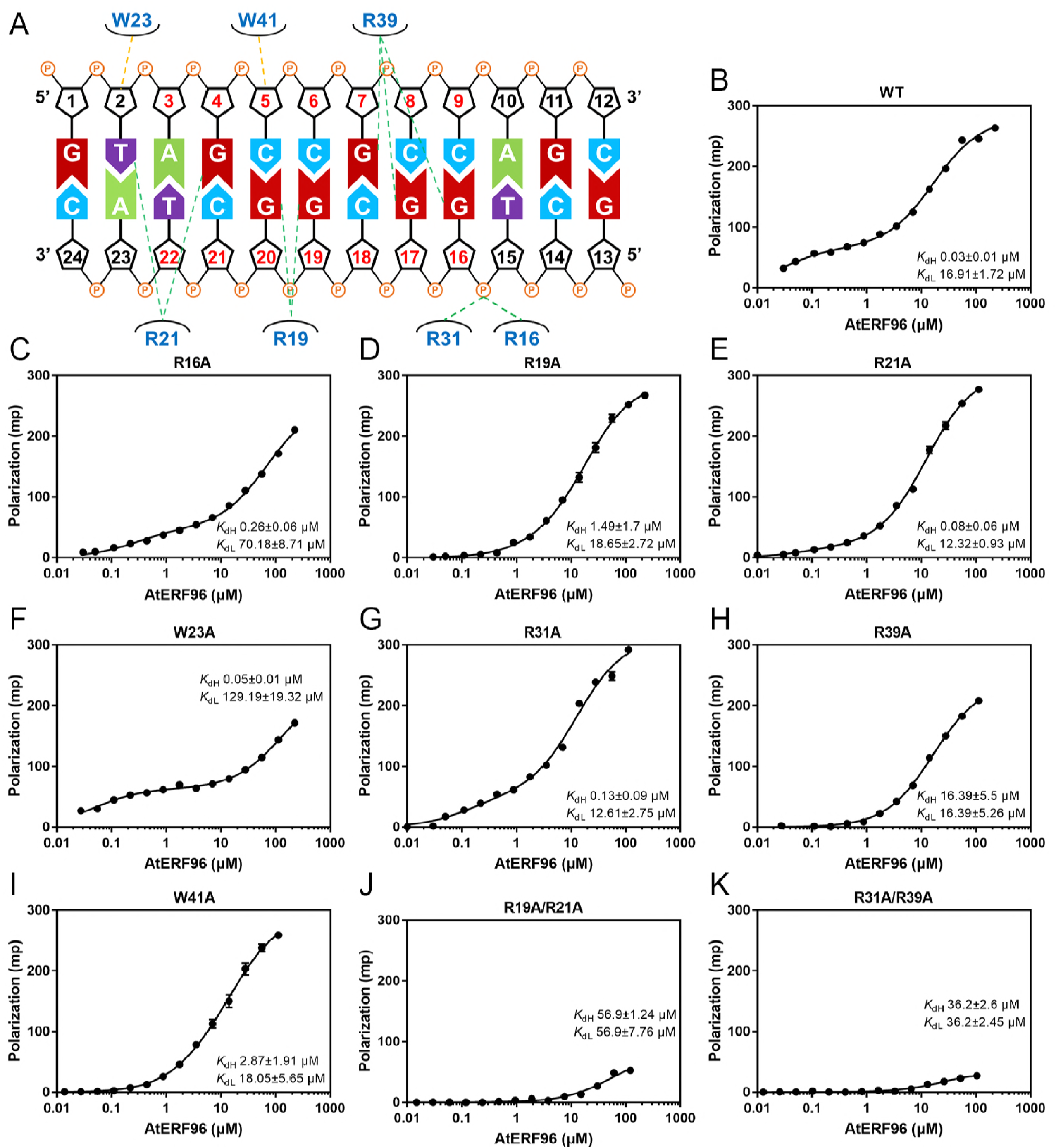
Fluorescence polarization analysis in the AP2/ERF domain of the AtERF96 protein. (**A**) Schematic diagram of the interaction network between AtERF96 protein and GCC12 DNA probe. The critical residues for protein-DNA interaction at the AP2/ERF domain are indicated. (**B-K**) Binding curves of the AtERF96 wild-type (**B**), R16A (**C**), R19A (**D**), R21A (**E**), W23A (**F**), R31A (**G**), R39A (**H**), W41A (**I**), R19A/R21A (**J**), and R31A/R39A (**K**) proteins with the GCC12 DNA probes. All data are representative of three independent experiments with the two-site binding equation, and the error is calculated as the standard deviation.

**Figure 4.**
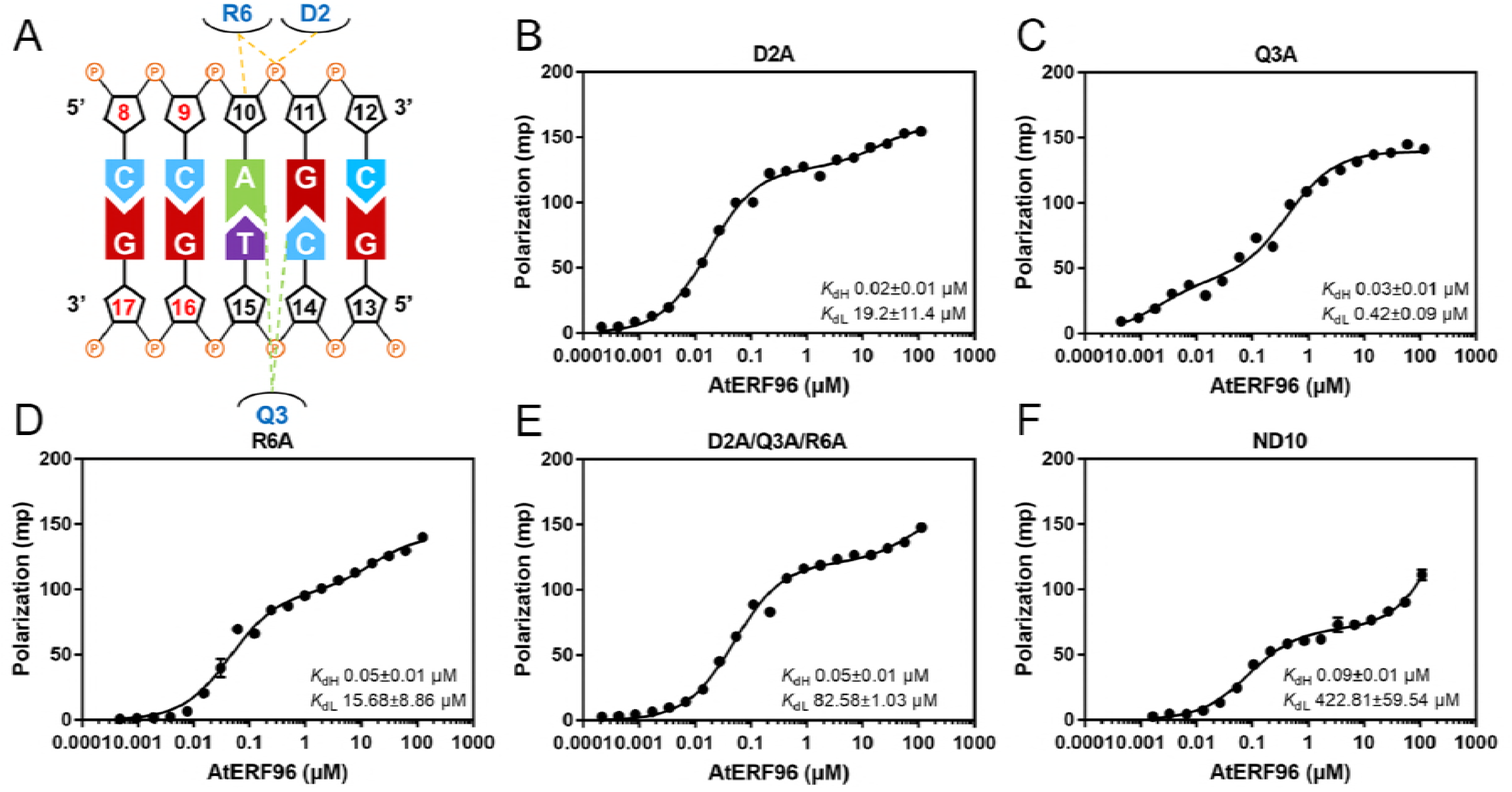
Fluorescence polarization analysis in the N-terminus of the AtERF96 protein. (**A**) Schematic diagram of the residues-nucleotides binding in the N-terminus of AtERF96 are shown. (**B-F**) Binding curves of the AtERF96 D2A (**B**), Q3A (**C**), R6A (**D**), D2A/Q3A/R6A (**E**), and the N-terminal truncation (ND10) (**F**) proteins with the GCC12 DNA probes. All data are representative of three independent experiments with the two-site binding equation, and the error is calculated as the standard deviation.

### Protoplast transactivation analysis

Our structural data showed that both the AP2/ERF domain and the N-terminal region of AtERF96 interact with the GCC box motif (Fig. 2E). We compared two GCC box motifs of the AtERF96–GCC11 and AtERF100–GCC11 complexes (Fig. 5A and B) and noticed that the α1 helix (M1-G9) of AtERF96 binds to the flanking region of the GCC11 DNA motif (Fig. 2E). The α1 helix contacts the template strand of the GCC box in the 5’ end region, resulting in a conformational change of the template strand and slight flipping of the ten nucleotide base pairs from C7 to G16 (Fig. 5C and D). In view of the N-terminal region of AtERF96 altering the DNA architecture of GCC11 in the crystals, we performed a transactivation analysis in AtERF96-overexpressing protoplasts to investigate whether the N-terminal region of AtERF96 is involved in transcription regulation. A luciferase (LUC)-encoding reporter gene, *PDF1.2 pro:LUC*, which contains two copies of the GCC box sequence from the *PDF1.2* promoter, and an effector plasmid consisting of each *AtERF* under the control of the cauliflower mosaic virus (CaMV) 35S promoter, were co-infiltrated into the *Arabidopsis* protoplasts (Fig. 6A). The results showed that LUC activity was not affected by the N-terminal-mutated or N-terminal-truncated AtERF96 proteins. However, decreased LUC activity was detected when the reporter plasmids were coexpressed with the effector plasmids of the AtERF96 R6A mutant (Fig. 6B). The data indicate that the N-terminal region of AtERF96 has a minor effect on transcription of the target gene. By contrast, the LUC activities affected by the AP2/ERF domain mutants of the AtERF96 effector plasmids were significantly reduced (Fig. 6C). These data coincide with the results of the binding between the AtERF96 mutants and the GCC box probes from the FP analysis. Coexpression of the EDLL-truncated AtERF96 with the reporter construct resulted in significant LUC inactivation (Fig. 6C). The mutants W23A and W41A showed a reduced binding ability to the GCC box in the transactivation assay, consistent with the results of the FP analysis and fEMSA (Fig. 3, S5 and S6). Furthermore, we observed two regions of the EDLL motif with high B-factor distributions, including the residues G80 to S84 and V104 to Y109 (Fig. 6D) (Çevik, Kidd et al., 2012, Tiwari et al., 2012). The sequence alignment showed that glutamate, aspartate, and leucine are enriched and highly conserved in the EDLL motif of group IX of the AP2/ERF family (Fig. 6E, Fig. S3). The results suggest that the EDLL motif is necessary for AtERF96 to interact with MED25, a subunit of the mediator complex in *Arabidopsis.*

**Figure 5.**
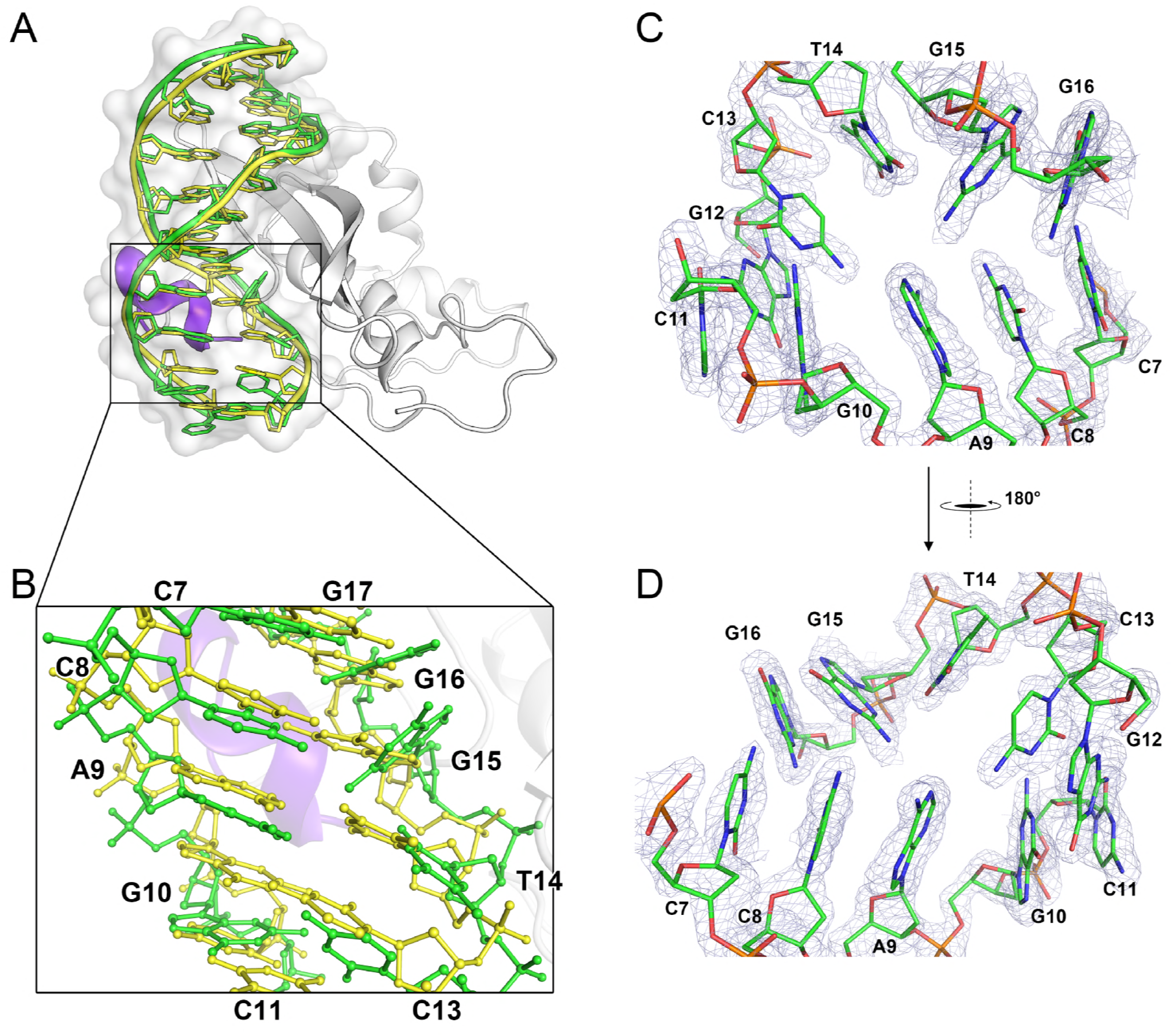
Conformational change of the DNA template strand by the binding of an N-terminal α1 helix in AtERF96. (**A**) Ribbon representation of the structural superposition of AtERF100-bound GCC11 DNA motif (yellow) and AtERF96-bound GCC11 DNA motif (green). (**B**) Zoom-in view of the base pairs of the superimposed AtERF100-bound GCC11 DNA motif (yellow) and AtERF96-bound GCC11 DNA motif (green) are shown as sticks. (**C and D**) Electron density map at the 5’ end of template strand in the AtERF96-bound GCC box motif. Minor groove (**C**) and major groove (**D**) views of the GCC11 DNA motif with nucleotides C7 to G16 from the structure of AtERF96–GCC11 complex are contoured at the 1.5 *σ* of 2 *F*_o_–*F*_c_ map.

**Figure 6.**
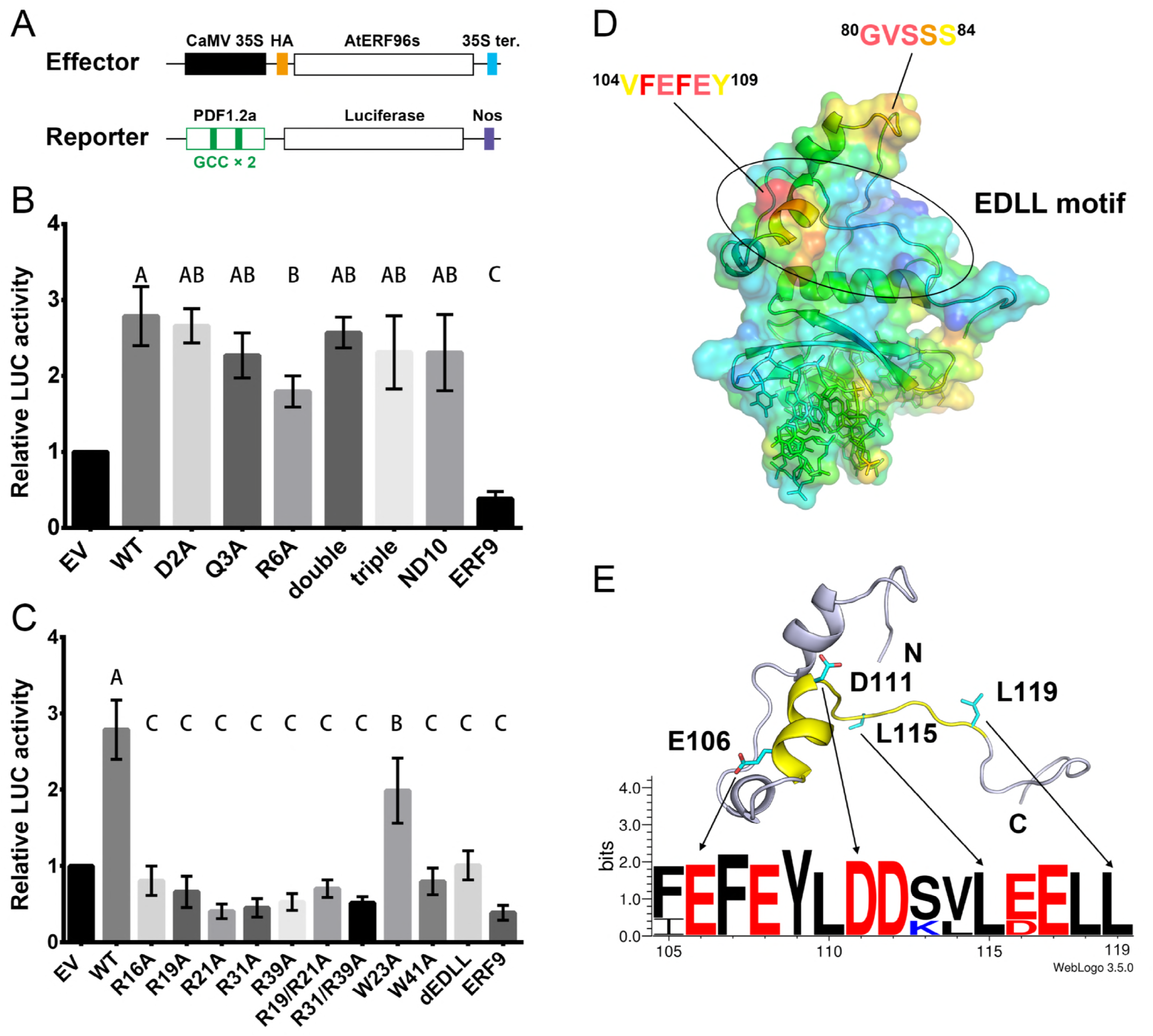
Transient expression assay and EDLL motif of AtERF96. (**A**) Schematic diagram of the reporter and effector plasmids used in transient assays. Effector plasmids were under the control of the cauliflower mosaic virus (CaMV) 35S promoter. The plasmids constructed with the Arabidopsis *PDF1.2a* promoter, which contains the two GCC boxes, were fused to a firefly luciferase gene as the reporter. HA tag, human influenza hemagglutinin tag. 35S ter, CaMV 35S terminator. Nos, the terminator signal of the gene for nopaline synthase. (**B and C**) Relative LUC activity from transient expression analysis of *PDF1.2a* promoter co-infiltrated with a plasmid containing AtERF96 genes fused to the 35S promoter. Plots of the LUC activity level influenced by AtERF96 genes with the mutations of N-terminal region (**B**) or AP2/ERF domain region (**C**) are shown, and the ERF9 is a negative control. EV, empty vector; WT, wild type; double, D2A/Q3A double mutations; triple, D2A/Q3A/R6A triple mutations; ND10, N-terminal deletion of first 10 residues; dEDLL, C-terminal deletion of the residues R102 to K131. Multiple comparisons of group vectors were performed using Fisher’s least-significant-difference (LSD) procedure. (**D**) Crystallographic B-factor distribution of the AtERF96–GCC11 complex. The residues of relatively higher B-factor are highlighted from high to low values as red > orange > yellow colors. (**E**) Ribbon representation and sequence logo of the AtERF96 EDLL motif. The designated EDLL region and conserved residues are indicated as yellow ribbons and cyan sticks. Sequence logo of the EDLL motif is shown through the full-length alignment of the paralogues from AtERF95 to AtERF98. The bit score indicates the information content for each position in the sequence.

### DNA binding specificity of AtERF96

To determine whether AtERF96 proteins interact with non-GCC box motifs, we tested three DNA motifs with GC-rich sequences: P box, CS1 box, and DRE box (A/GCCGAC) (Hao, Yamasaki et al., 2002). We designed these three probes with fluorescein fused to the 5’- or 3’-end and tested the binding ability between AtERF96 proteins and these DNA motifs using fEMSA and FP analyses. The GCC12 probe and the W box (TTGACC) probe were used as positive and negative controls for fEMSA analysis, respectively. (Fig. S7B). The fEMSA results showed that AtERF96 protein has a slight binding ability to P box, CS1 box, and DRE box motifs (Fig. 7A). The FP assay also revealed that the K_dHi_ levels of these motifs to AtERF96 protein were much weaker than the GCC box by 8 to 25 fold, respectively (Fig. 7B–D, Table S8). By contrast, the influence of the K_dLo_ levels on the P box and DRE box motifs was insignificant (Table S8). These data suggest that the AtERF96 protein retains a limited binding ability for other DNA motifs through non-specific interactions.

**Figure 7.**
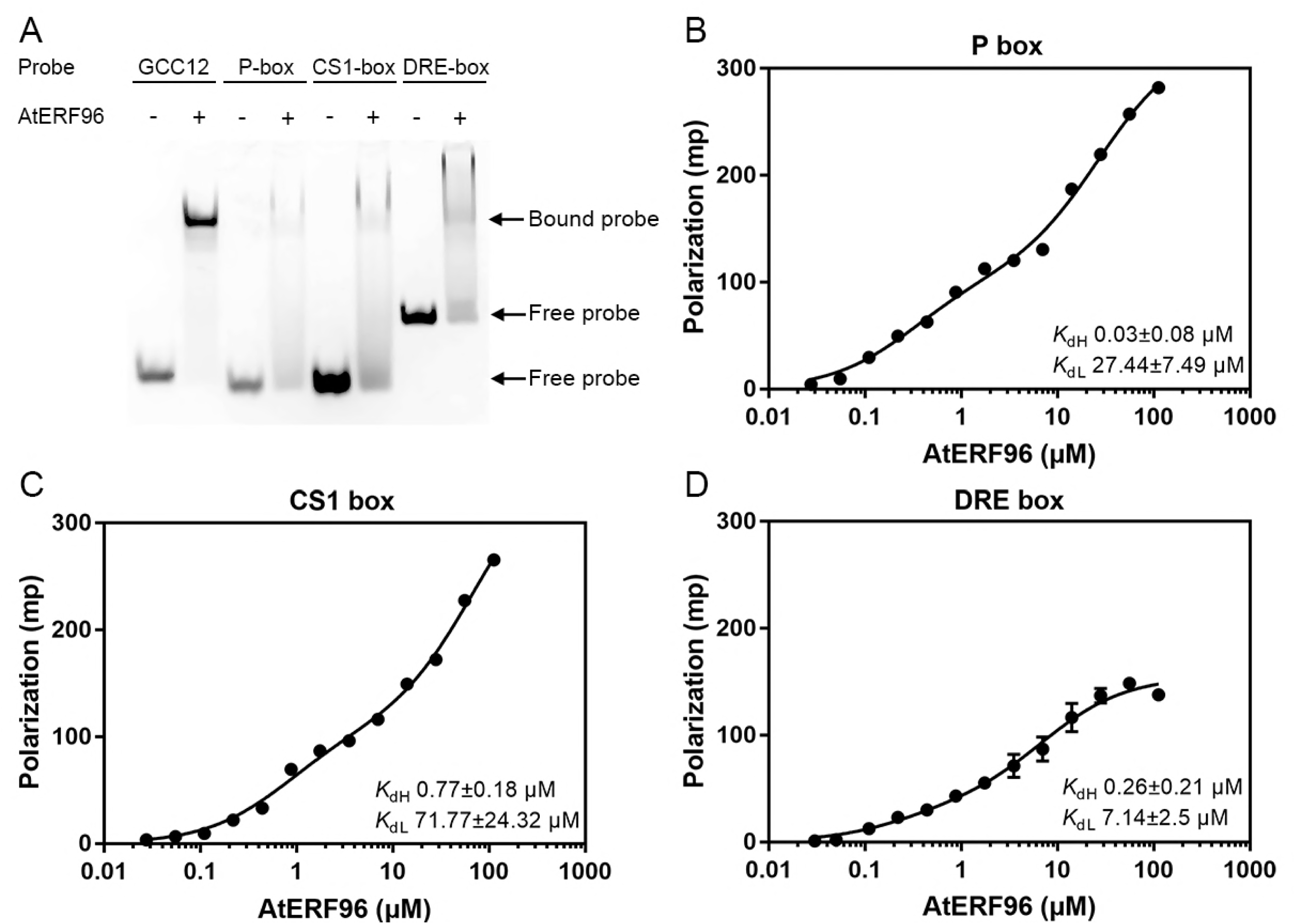
Characterization of binding ability between AtERF96 proteins and various DNA motifs. (**A**) EMSA binding analysis of AtERF96 proteins with the GCC12, P box, CS1 box, and DRE box probes. (**B-D**) Fluorescence polarization analysis of AtERF96 proteins with the P box (**B**), CS1 box (**C**), and DRE box (**D**) DNA probes. All data are representative of three independent experiments with the two-site binding equation, and the error is calculated as the standard deviation.

## DISCUSSION

AP2/ERF-family proteins regulate transcription by recognizing the GCC box sequence in the promoters of target genes (Mizoi, Shinozaki et al., 2012a). Group IX of the AP2/ERF family is composed of three subgroups, IX-a, IX-b, and IX-c, characterized by the conserved motifs (CM) CMIX-3, CMIX-2, and CMIX-1 (specifically, the EDLL motif), respectively (Nakano et al., 2006). Among these, the members of group IX of AP2/ERFs have been linked to defensive gene expression in response to pathogen infection (Berrocal-Lobo et al., 2002, Gu, Wildermuth et al., 2002, Gutterson & Reuber, 2004). AtERF96 phylogenetically belongs to group IX-c of the AP2/ERF gene family. The amino acid sequence of AtERF96 is similar to that of AtERF95, AtERF97, and AtERF98, and is relatively smaller than that of the other members of group IX. Recently, Catinot and colleagues showed that overexpressed *AtERF96* enhances *Arabidopsis* resistance to necrotrophic pathogens, such as the fungus *Botrytis cinerea* and the bacterium *Pectobacterium carotovorum* (Catinot et al., 2015). A microarray assay coupled to chromatin immunoprecipitation-PCR of overexpressed AtERF96 revealed that AtERF96 regulates the activation of JA/ET-responsive genes, such as *PDF1.2a*, *PR*-*3*, and *PR*-*4*, as well as the transcription factor ORA59, through direct binding to existing GCC elements in their promoters (Berrocal-Lobo et al., 2002, Catinot et al., 2015, Pre et al., 2008).

In this study, we determined the crystal structure of the AtERF96–GCC11 complex, including an AP2/ERF domain and an EDLL motif at a resolution of 1.76 Å (Fig. 1C, Table 1). The conformation of the AP2/ERF domain in AtERF96 shows a similar framework to AtERF100 upon binding to the target DNA (Fig. S2) (Allen et al., 1998). Nevertheless, the potential propensity of residue-nucleotide interactions shows some differences between these two structures. For example, residue R31 of AtERF96 (R162 of AtERF100) contacts the phosphate group of nucleotide G15; at the same structural position, the arginine of AtERF100 binds to the guanine base. Residue R21 of AtERF96 (R152 of AtERF100) contacts nucleotides T1 and G3 at the 5’-end, but residue R152 of AtERF100 binds to nucleotide G19 closer to the 3’-end of another strand. Residue R39 of AtERF96 (R170 of AtERF100) contacts three guanines, G6, G15, and G16, instead of the sugar-phosphate backbone of nucleotide C5. There are two causes for these differences: one is the discrepancy of the polar residues from the few non-conserved amino acids between these two AP2/ERF domains; the other is the influence of the neighboring α1 helix at the N-terminus of AtERF96. The N-terminal α1 helix interacts with the flanking region following the core sequence of the GCC11 motif at the minor groove. Interestingly, residue Q3 of the α1 helix provides polar interactions with nearby nucleotides, especially G10 and T14, resulting in unpairing and unstacking of base pairs from C7 to G16 (Fig. 5C and D). We used the 3DNA suite of programs to analyze the conformation of DNA base pairs in the residue-binding region. Results indicated that nucleotides C8, T14, and G15 exhibit shifting, tilting, and rolling (Table S4). Disruption of base stacking in single-stranded polynucleotides significantly alters the base pair conformation, leading to a lack of information on the spatial configurations of base pairs C8-G15 and A9-T14 (Table S5 and S6). The residue-base interaction of R39-G16 combined with the shifting and twisting of base C7/C8 directly leads to a shear in the base pair C7-G16 (Table S5). Thus, the unpaired and unstacked nucleotides further affect the interaction with the residues of the β1–β2 strands at the major groove. This result explains why few conserved arginines in the AP2/ERF domain of AtERF96 show a binding mode distinct from AtERF100 using target nucleotides. No previous reports indicate that binding of the ethylene-responsive element binding factors with the GCC box motif results in the unstacking of DNA bases. Regarding AtERF96 acting as a positive regulator of target gene transcription, we suggest that binding of the N-terminal α1 helix with the 3’ flanking region may facilitate DNA unwinding for further transcription initiation.

Mutagenesis coupled with binding experiments confirmed the relevance of the protein-DNA contacts identified in our structure and helped us delineate the residue conservation in both the AP2/ERF proteins and the GCC box DNA targets. We analyzed the binding efficiency of AtERF96 wild-type and various mutants via fluorescence polarization analysis. We noticed that the interaction of AtERF96-GCC box showed the capacity for both specific and non-specific binding in our FP analysis. The polarization curve of wild-type AtERF96 showed a clear trend with two rises, which can be also observed in the results of R16A, W23A, and R31A mutants. (Fig. 3B, C, F and G). Interestingly, when attempting to analyze the data using the one-site binding equation: *y* = B_max_ × *x*/K_dHi_ + *x*, the data of wild-type and mutants again were difficult to fit to the sigmoid curve (Fig. S4). However, the equation can fit the data of R19A, R21A, R39A, W41A, and double mutants (Fig. S4). The results imply that some residues are crucial for the recognition of the GCC box, and that the mutations caused the functional loss of specific binding. Among these, R19 and R39 showed the major influence in the specific binding, due to their interactions with the bases G7, G16, G17, G19 and G20 in the GCC12 probe (G6, G15, G16, G18 and G19 in the GCC11 probe) (Fig. 3, Table S7). At the N-terminus of AtERF96, all mutants retained the two-site binding feature in the raw data (Fig. 4). The ND10 truncated protein showed a limited effect on specific binding, accompanied by a raised K_dHi_ level compared to wild-type, suggesting that the N-terminal region is not involved in GCC box recognition (Fig. 4F). In view of the above, we suggest that the K_dHi_ is implicated in the residue-base conservation, whereas the K_dLo_ reflects the stabilization of residue-sugar phosphate backbone, according to the effects caused by mutations of conserved residues of their specific functions. To better understand the impact of AtERF96 mutations *in vivo*, we designed a transactivation analysis in *Arabidopsis* protoplasts with an overexpressing effector and a luciferase-fused reporter. Although the N-terminal α1 helix of AtERF96 made contact with the 5’ end of the template strand and structurally disrupted DNA base pairing, the N-terminal mutants only showed limited influence on the transactivation analysis (Fig. 5B). These results show that the α1 helix acts as an auxiliary domain in promoting transcription initiation. The transactivation assay revealed that most mutations of conserved arginines in the AP2/ERF domain seriously disrupted protein-DNA interactions, including the conserved tryptophans W23 and W41 (Fig. 6C). However, we noticed that the sequence region excluding the AP2/ERF domain is highly diverse in all of group IX members in the AP2/ERF family, meaning that the N-terminal region of other ERFs are structurally distinct from AtERF96 (Fig. S3). On the other hand, the EDLL-truncated AtERF96 lost its transactivation function *in vivo*, suggesting that the EDLL motif indeed interacts with the MEDIATOR25 subunit of the eukaryotic Mediator complex (Çevik et al., 2012). Previous work confirmed that MED25 interacts with the four members of group IX of the AP2/ERF family, i.e., AtERF92, AtERF93, ORA59, and TDR1/AtERF98. Interestingly, the α5 helix of the EDLL motif exhibited a significantly increased B-factor in the whole structural data, implying that this region probably plays an important role in attaching to the MED25 subunit (Fig. 6D). Nevertheless, structural studies and mechanistic insights into MED25 are still needed.

In summary, we have shown that AtERF96, an AP2/ERF-family regulator recognizing the GCC box DNA motif, is an ethylene-responsive transcription factor that directly modulates the defense-related gene *PDF1.2a.* Our studies of the AtERF96–GCC11 complex provide a structural framework for AP2/ERF transcription factors, including the binding capability of the AP2/ERF domain, together with the influence of DNA base-pair opening via the N-terminal helix of AtERF96.

## SUPPLEMENTARY DATA

Supplementary Data are available at The EMBO Journal Online.

## ACKNOWLEDGEMENTS

We thank Dr. Laurent Zimmerli for providing the material of *AtERF96* gene construct. We thank Uni-edit (www.uni-edit.net) for editing and proofreading this manuscript. We thank the technical supports provided by the “Synchrotron Radiation Protein Crystallography Facility of the National Core Facility Program for Biotechnology, Ministry of Science and Technology” and the “National Synchrotron Radiation Research Center”, a national user facility supported by the Ministry of Science and Technology, Taiwan. This work is partly supported by Technology Commons, College of Life Science, National Taiwan University, Taiwan.

## FUNDING

Financial supports were from the Taiwan Ministry of Science and Technology (MOST 106-2313-B-002-005) and National Taiwan University (NTUCCP-106R891508) to Yi-Sheng Cheng.

## Conflict of interest statement

None declared.

